# Bayesian estimation of trunk-leg coordination during walking using phase oscillator models

**DOI:** 10.1101/2024.04.25.591041

**Authors:** Haruma Furukawa, Takahiro Arai, Tetsuro Funato, Shinya Aoi, Toshio Aoyagi

## Abstract

In human walking, the legs and other body parts coordinate to produce a rhythm with appropriate phase relationships. From the point of view for rehabilitating gait disorders, such as Parkinson Disorders, it is important to understand the control mechanism of the gait rhythm. A previous study showed that the antiphase relationship of the two legs during walking is not strictly controlled using the reduction of the motion of the legs during walking to coupled phase oscillators. However, the control mechanisms other than those of the legs remains unknown. In particular, the trunk moves in tandem with the legs and must play an important role in stabilizing walking because it is above the legs and accounts for more than half of the mass of the human body. This study aims to uncover the control mechanism of the leg-trunk coordination in the sagittal plane using the coupled phase oscillators model and Bayesian estimation. We demonstrate that the leg-trunk coordination is not strictly controlled, as well as the interleg coordination.

## 1 Introduction

Trunk control is crucial for achieving a stable gait. The trunk accounts for over 50% of the total body weight (Dempster, 1955; Plagenhoef et al., 1983), and a slight trunk tilt can produce significant moments at each joint (Preece and Alghamdi, 2021). Additionally, due to the narrower Base of Support (BoS) in humans compared to quadrupedal animals, there is a tendency for the center of mass to protrude beyond the perimeter of the BoS (Winter et al., 1990). Moreover, the center of mass during walking is higher than that of the legs, which makes walking mechanically unstable.

Numerous researchers have studied the control of legs during walking (Bauby and Kuo, 2000; Collins and Kuo, 2013; Arvin et al., 2016). These studies are frequently discussed in terms of stability of the mechanical system. It has been reported that stability in the frontal plane is strongly related to step width (Bruijn and van Dieën, 2018). The leg motion in the sagittal plane is more stable than that in the frontal plane (Bauby and Kuo, 2000; McGeer, 1990).

This study focuses on the stability of gait rhythm, rather than the mechanical stability described above. In walking, each part of the body has its own rhythm. These rhythms interact with each other and create appropriate phase relationships. For instance, the right and left legs move alternately (Alexander, 2002), and the arms move in tandem with them. This rhythmic coordination is typically stable in healthy young people, whereas it can be disrupted in the elderly and in patients with Parkinson’s disease (Plotnik et al., 2007, 2008). To assist those with gait disorders, it may be useful to uncover the mechanism of rhythmic coordination.

There are few studies with regard to the stability of gait rhythm, despite its significance. A previous study suggests that the antiphase relation of the left and right legs is importatnt to stabilize walking, but it is not strict (Arai et al., 2023). That is, interleg coordination is not tightly controlled. The trunk also moves in tandem with the legs and trunk-leg coordination must play an important role in stabilizing walking.

However, the trunk-leg coordination during walking remains largely unclear.

The aim of this study is to clarify the mechanisms of trunk-leg coordination in the sagittal plane during walking from the point of view of the rhythm control. To achieve this, we use nonlinear dynamical systems and Bayesian inference method. We model the rhythmic motion of the legs and the trunk with coupled phase oscillators, which is based on the phase reduction theory (Kuramoto, 1984; Winfree, 1980; Nakao, 2016). Then, we identify the model parameters from the time-series data using Bayesian inference method (Ota and Aoyagi, 2014).

## 2 Phase oscillator models

The trunk and legs show complex behavior during walking, but each of them has a rhythm. These rhythms interact with each other. In this study, a coupled phase oscillators model that renormalizes the complex interactions of muscles and skeleton comprehensively is used for the trunk and legs to elucidate the control mechanisms of the rhythm in the sagittal plane. This model focuses only on rhythm among the higher-dimensional behaviors.

Following the phase reduction theory, if each element composing a system has a rhythm generated by a limit cycle, the rhythm is represented by a phase variable (Kuramoto, 1984; Winfree, 1980; Nakao, 2016). From this result, we model the motion of the right and left legs and the trunk by coupled limit cycle oscillators and we use *ϕ*_R_(*t*), *ϕ*_L_(*t*), *ϕ*_T_(*t*) to represent these phases respectively (see Figure 1(a)). These phases are supposed to monotonically increase if a noise and an external perturbation are not added to the oscillators. These increase rates are called natural frequencies, denoted by *ω*_R_, *ω*_L_, *ω*_T_.

**Figure 1:**
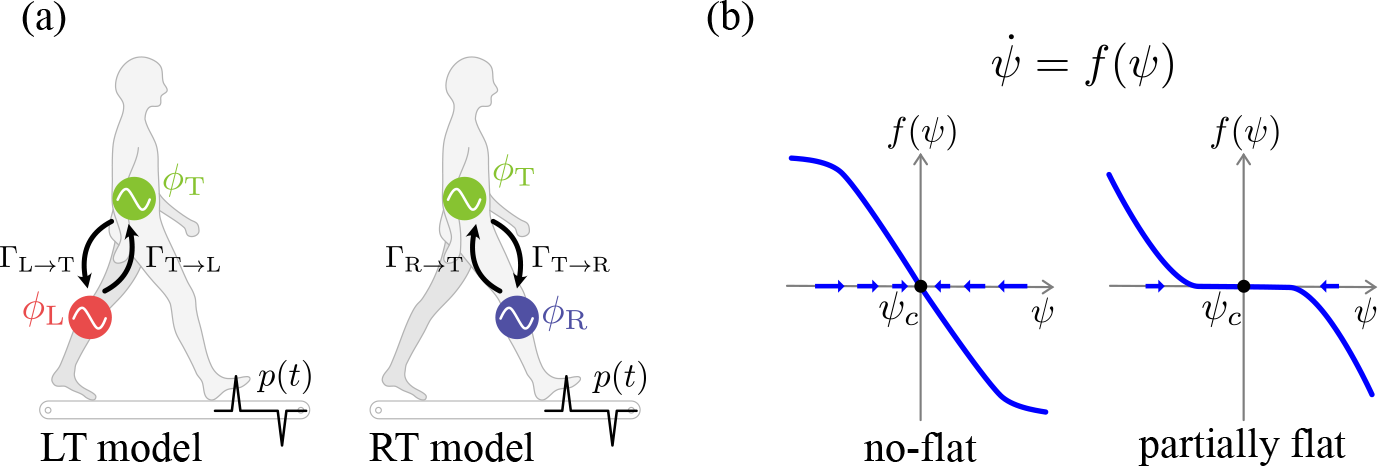
(a) Two coupled phase oscillators models proposed in this study. The legs and trunk are assumed to have a single rhythm during gait, and one oscillator is assigned to each of them; the two phase oscillators are subject to external perturbations and stochastic fluctuations. (b) A correspondence relationship between the shape of the spontaneous dynamics *f* (*ψ*) of the relative phase *ψ* and its control mechanism. Suppose that *ψ* is locked at *ψ* = *ψ*_*c*_. If *f* (*ψ*) is not flat near *ψ*_*c*_ as shown at the left, then even a slight shift of *ψ* is corrected sensitively. By contrast, if *f* (*ψ*) has a flat interval in the vicinity of *ψ* = *ψ*_*c*_ as shown at the right, then the control of *ψ* is not actively performed until it shifts to a certain value.

Trunk and leg behaviors in the sagittal plane are synchronized in a 1:2 ratio (see Figure 2(b)). With this consideration, we model the motion of the legs and the trunk as follows:

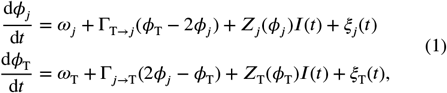

where *j* ∈ {R, L}, Γ_*j*→*k*_ is a phase coupling function that represents the action from the oscillator *j* to *k. Z*_*j*_ (*ϕ*_*j*_) is a phase sensitivity function of the oscillator *j*, i.e., the linear response to an external perturbation *I*(*t*). *ξ*_*j*_ (*t*) denotes an independent Gaussian white noise that satisfies

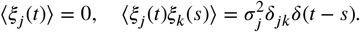

Here, *δ*_*jk*_ is Kronecker’s delta and *δ*(*t* − *s*) is Dirac’s delta function. Henceforth, we call the models RT-model for *j* = R and LT-model for *j* = L.

**Figure 2:**
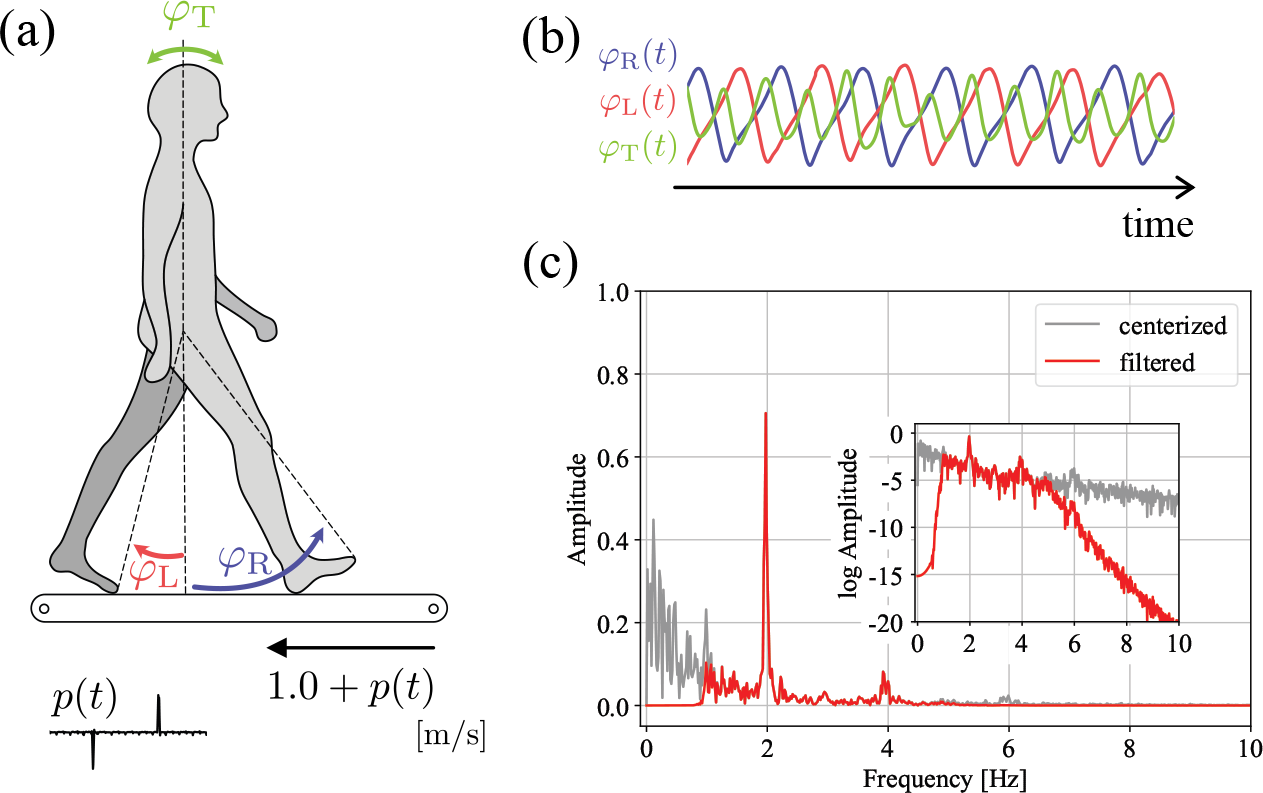
(a)The definitions of angles used to reconstruct the phase time series. (b)Normalized angular time series.(c)Changes in the frequency spectrum before applying the filter (gray) and after (red).

The stability of phase synchronization is determined by the spontaneous dynamics of the phase oscillator model. We define the relative phase between the right leg and the trunk as *ψ*_RT_ := 2*ϕ*_R_ − *ϕ*_T_, that between the left leg and the trunk as *ψ*_LT_ := 2*ϕ*_L_ − *ϕ*_T_.

Its time evolution of *ψ*_RT_ and *ψ*_LT_ without the external perturbation is represented as follows:

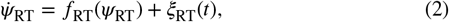

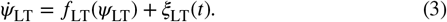

where

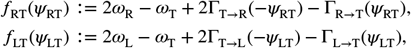

and the noise *ξ*_RT_(*t*), *ξ*_LT_(*t*) satisfies

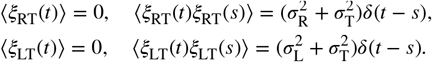

The shapes of the funtions *f*_RT_(*ψ*_RT_), *f*_LT_(*ψ*_LT_) imply the control mechanism of the rhythm. Considering the stability of the gait rhythm in healthy individuals, it is expected that they sensitively correct small shifts in the relative phase. In this case, the function intersects the axis transversely at one point, as illustrated on the left side of Figure 1(b). If the control of rhythm is not strict, by contrast, the function intersects the axis tangentially, as illustrated on the right side of Figure 1(b), with a flat interval in the vicinity of the intersection point. Some studies suggest that the latter control mechanism is observed during bipedal walking (Arai et al., 2023) and upright posture (Asai et al., 2009).

Note that the motion of the left and right legs is modeled with

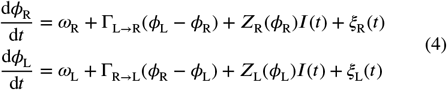

in previous research (Arai et al., 2023), which we call this RL-model. We follow this approach.

## 3 Methods

### 3.1 measuring

All subjects (A–H) were healthy men (*n* = 8; age 21– 23 years; height 161–182 cm, weight 50.2–79.5 kg). The subjects walked on a treadmill (ITR3017, BERTEC corporation) while we captured three-dimensional coordinates using a motion capture system (MAC3D Digital RealTime System; NAC Image Technology, Inc.). The data sampling rate was 500 Hz. The subjects outfitted reflective markers on various regions of their bodies: the head, the top of the acromion, the greater trochanter, the lateral condyle of the knee, the lateral malleolus, the head of the second metatarsal, and the heel. These markers were affixed at one location on both the left and right sides.

In human walking, the legs and the trunk exhibit a stable synchronization; therefore, if no perturbations are applied, the experimental data tends to be localized in only a small portion of the data space. To solve this problem, we introduced perturbations through the change of the velocity of the belt during the experiment to disrupt the synchronization.

The unperturbed belt speed was set at 1.0 m/s. We applied acceleration or deceleration perturbation to the speed from this point. For acceleration perturbations, the speed was increased to 1.6 m/s over 0.1 seconds, and then returned to 1.0 m/s over the next 0.1 seconds. For deceleration perturbations, the speed was decreased to 0.4 m/s over 0.1 seconds, and then returned to 1.0 m/s over the next 0.1 seconds. Whether to apply only acceleration perturbations (acceleration condition), only deceleration perturbations (deceleration condition), or a random combination of the two (mixed condition) varied across trials.

Each trial of the experiment lasted approximately 60 seconds. The first 10 seconds of each trial involved no perturbations. Subsequently, perturbations were applied every 5 seconds, with a total of 10 perturbations per trial. Subject A performed 25 trials with acceleration condition and 24 trials with deceleration condition. Subject B performed 25 trials with acceleration condition and 25 trials with deceleration condition. Subjects C–H performed 15 trials each with acceleration condition, deceleration conditions, and mixed conditions.

This study was approved by the Ethics Committee of Doshisha University. Written informed consent was obtained from all subjects. We use a part of the measurement data from previous studies (Funato et al., 2015, 2016) for different purposes. Markers utilized to construct the time series of the trunk in this study are not used in these studies.

### 3.2 data processing

Firstly, the angular time series of the legs and the trunk in the sagittal plane were calculated from the captured three dimensional coordinates, as shown in Figure 2(a). With regards to the legs, the sagittal plane projection of the line connecting the greater trochanter and the head of the second metatarsal was computed, along with the angle it forms with the vertical direction, which we call the leg axis angle. The right leg axis angle is denoted by *φ*_R_(*t*) and the left by *φ*_L_(*t*). With regards to the trunk, the angle formed between the sagittal plane projection of the line connecting the midpoint of the acromia and the greater trochanters on both sides and the vertical direction was calculated, which we call the trunk flexion angle.

While both of the leg axis angles exhibited distinct rhythms in all subjects, the trunk flexion angle tended to less variable and varied more among individuals. To reconstruct the phase time series of trunk effectively, a frequency bandpass filter (from 1.0 Hz to 5.0 Hz) was applied to the centerized time series to remove trends and noise. The time series data of the trunk flexion angle after filtering were denoted as *φ*_T_(*t*). An example of the time series of a subject G after filtering is shown in Figure 2(b), and the frequency spectrum is shown in Figure 2(c).

#### 3.2.1 reconstruction of phase time series

To reconstruct the phase time series *ϕ*_*j*_ (*t*), the angular time series *φ*_*j*_ (*t*) was projected onto the complex plane, yielding the argument *θ*_*j*_ (*t*) (Strogatz, 2003),

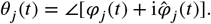

Here, 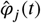 is the Hilbert transform of *φ*_*j*_ (*t*) and 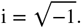.

While *θ*_*j*_ (*t*) ∈ [0, 2*π*) for all *t*, this is not suitable for the phase oscillator model because in general *θ*_*j*_ (*t*) does not satisfy the condition expected of the phase, 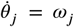 with no perturbations. Hence, following the method proposed by Kralemann (Kralemann et al., 2007, 2008), *ϕ*_*j*_ (*t*) was obtained as

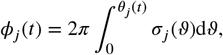

where *σ*_*j*_ (*ϑ*) is a probability density function of *θ*_*j*_ (*t*) constructed from the time-series data. This probability density function is constructed from time-series data obtained from all trials of the experiments, for each subject and each perturbation condition. We used this *ϕ*_*j*_ (*t*) as the phase of the oscillator.

Furthermore, perturbation *p*(*t*) was rescaled to *I*(*t*) as follows:

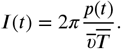

Here, 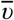 is the average speed of the treadmill belt, and 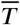 is the average walking period.

The time series of phases and relative phases obtained through the aforementioned procedure are illustrated in Figure 3.

**Figure 3:**
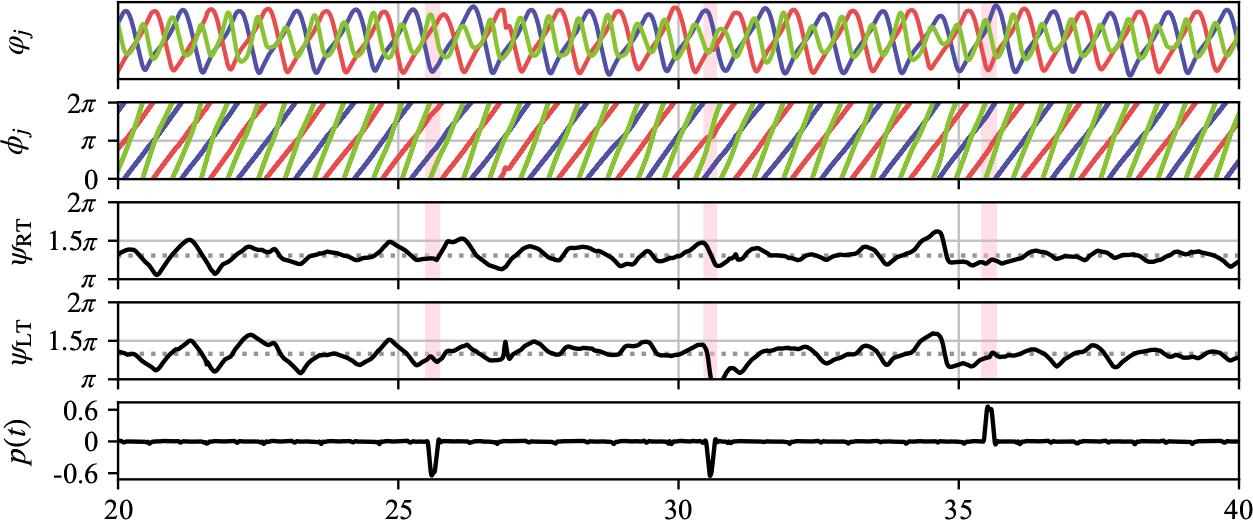
The observed time series and the constructed phase time series from them. The colors red, blue and green correspond to the time series with regard to the left leg, the right leg and the trunk respectively. The segment at the bottom indicates an external perturbation. The pink band indicates the times when external perturbations are present.

#### 3.2.2 the derivative of the phase and test for outliers

The reconstructed phase time series is, in fact, a discrete time series of length *T* sampled at a certain rate Δ*t*, rather than continuous in time. Therefore, we denote the observed phase time series of an oscillator *j* as 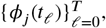, and the corresponding perturbation time series at each time step as 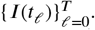. Phase derivatives (phase velocity) 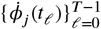 are approximated as follows:

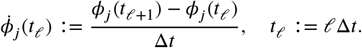

To statistically exclude outliers in the derivatives due to noise, we employ the Smirnov-Grubbs test (*p* < 0.1). During the test, timestamps with perturbations are excluded.

### 3.3 model identification

Using the reconstructed time series, we identify the equations (1). Bayesian inference method proposed by Ota et al. (Ota and Aoyagi, 2014) is performed.

Since the coupling function and phase sensitivity are 2*π*-periodic, they can be expanded using Fourier series.

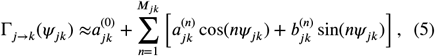

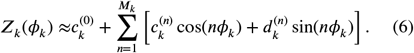

Here, 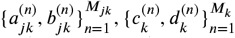 are coefficients of Fourier series and ***M***_*jk*_, ***M***_*k*_ are hyperparameters. Substituting them to the eq.(1), we obtain

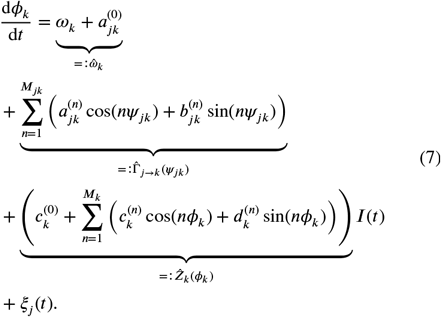

We identify these model parameters using the phase and perturbation time series.

To simplify the notation, we represent the Fourier coefficients collectively as follows:

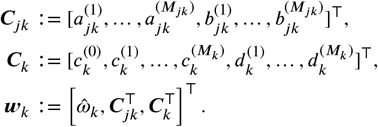

Unknown parameters in eq.(7) are ***w***_*k*_ and the variance of noise 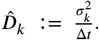. The estimation of them reduces to a linear regression, where the input variables are the phases of all oscillators and the external perturbation and the target variable is the phase velocity of oscillator *k*. Hence, following the Bayesian inference method (Ota and Aoyagi, 2014), we define a likelihood as

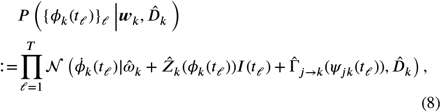

where 𝒩 denotes Gaussian distribution. The conjugated prior 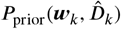 with respect to the likelihood (8) is Gaussian inverse-Gamma distribution,

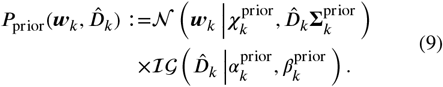

Here, 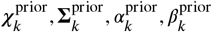 are hyperparameters of the conjugated prior, and inverse-Gamma distribution ℐ 𝒢 is defined

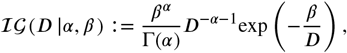

where Γ(*α*) is the Gamma function.

#### 3.3.1 updating hyperparameters

Due to the conjugacy of the prior, the posterior *P*_post_ is also Gaussian inverse-Gamma distribution with the parameters 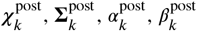. These can be expressed with the parameters of the prior distribution,

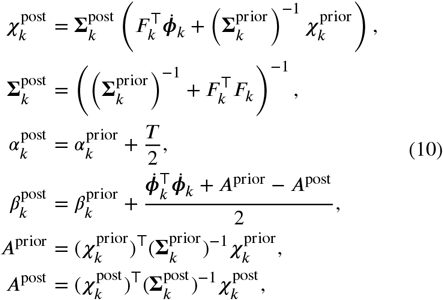

where *F*_*k*_ is the design matrix:

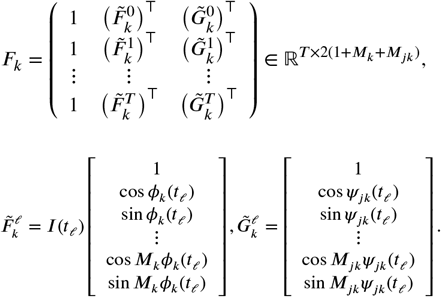

#### 3.3.2 model selection

The Fourier series of Γ_*j*→*k*_ (eq.(5)) and ***Z***_*k*_ (eq.(6)) are truncated at the order ***M***_*jk*_ and ***M***_*k*_, respectively. These hyperparameters represent the complexity of the model: the larger they are, the more complex the models can be intuitively interpreted as being. To select the appropriate ***M***_*jk*_ and ***M***_*k*_, we employ the evidence approximation. The model evidence *p*({*ϕ*_*k*_(*t*_*ℓ*_)}_*ℓ*_ |***M***_*μν*_, ***M***_*ν*_)is defined as

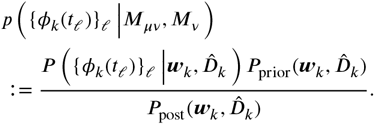

The most plausible order ***M***_*jk*_ and ***M***_*k*_ are set to maximize the model evidence.

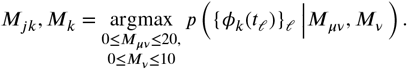

## 4 Result

### 4.1 phase sensitivity

First, we investigate the results regarding phase sensitivities. As shown in Figure 4(a), phase sensitivities of both legs *Z*_R_, *Z*_L_ are broadly divided into two regions: that close to zero and that taking positive values. The region close to zero corresponds to the swing phase, or the state where the foot is not in contact with the ground. These results are consistent with findings from the previous study (Arai et al., 2023).

**Figure 4:**
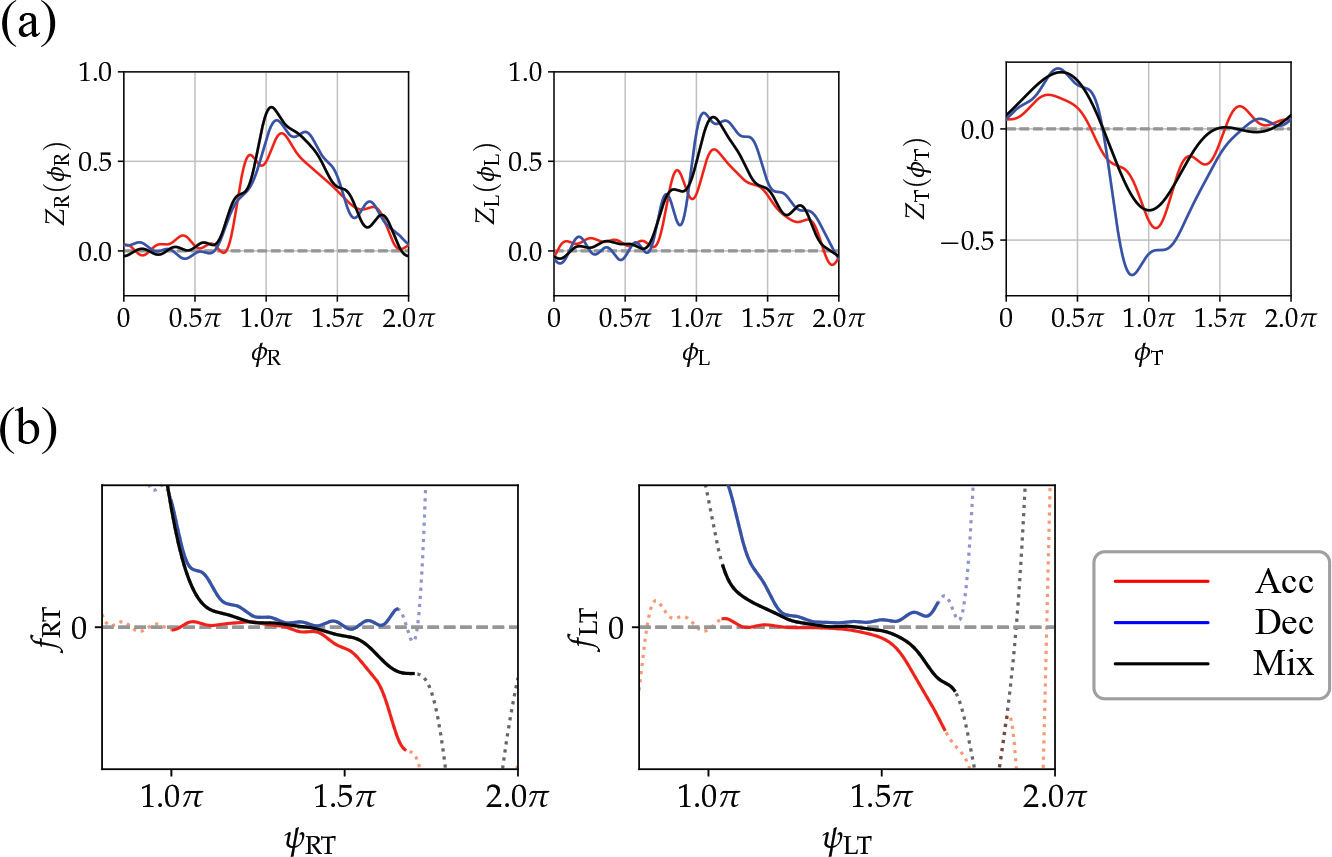
Results for the leg-trunk coordination for one subject (Subject G). Different colors represent patterns of different conditions: acceleration condition (Acc), deceleration condition (Dec), mixed condition (Mix). (a) the phase sensitivity of the left leg (left), the right leg (center), and the trunk (right). (b) the spontaneous dynamics of the relative phases, *ψ*_RT_ (left) and *ψ*_LT_ (right).

The phase sensitivity of the trunk *Z*_T_ differs from that of the legs. In the case that *ϕ*_T_ is in the vicinity of 0 to 0.7*π*, the phase response tends to be positive, while it becomes negative outside this range. The *ϕ*_T_ = 0.7*π* region corresponds approximately to when the trunk is most inclined forward, implying that avoiding falls is achieved with leaning the trunk forward. This observation aligns with experimental findings (Grabiner et al., 1993).

### 4.2 control of trunk-leg coordination

Next, we investigate the controls of trunk-leg coordination, *f*_RT_ and *f*_LT_ (Figure 4(b)). In both of them, flat intervals are observed. These flat intervals suggest that when the rhythm is perturbed, it remains uncorrected until the deviation surpasses a threshold. Similar results are obtained in interleg coordination in the previous research (Arai et al., 2023): the relative phase between the left and right legs is not strictly controlled and exhibits weak stability near *π*.

To quantitatively evaluate the flat intervals, we approximate the results using piecewise linear functions (Figure 5(a)). Hereafter, we analyze the results excluding the clearly exceptional subject (Subject C). Piecewise linear functions are fitted using least squares regression with four parameters: the *y*-coordinates (*y*_1_, *y*_2_) of the endpoints of the data range and the *x*-coordinates (*x*_1_, *x*_2_) of the endpoints of the flat interval, yielding the length of the flat interval, *x*_2_ − *x*_1_.

**Figure 5:**
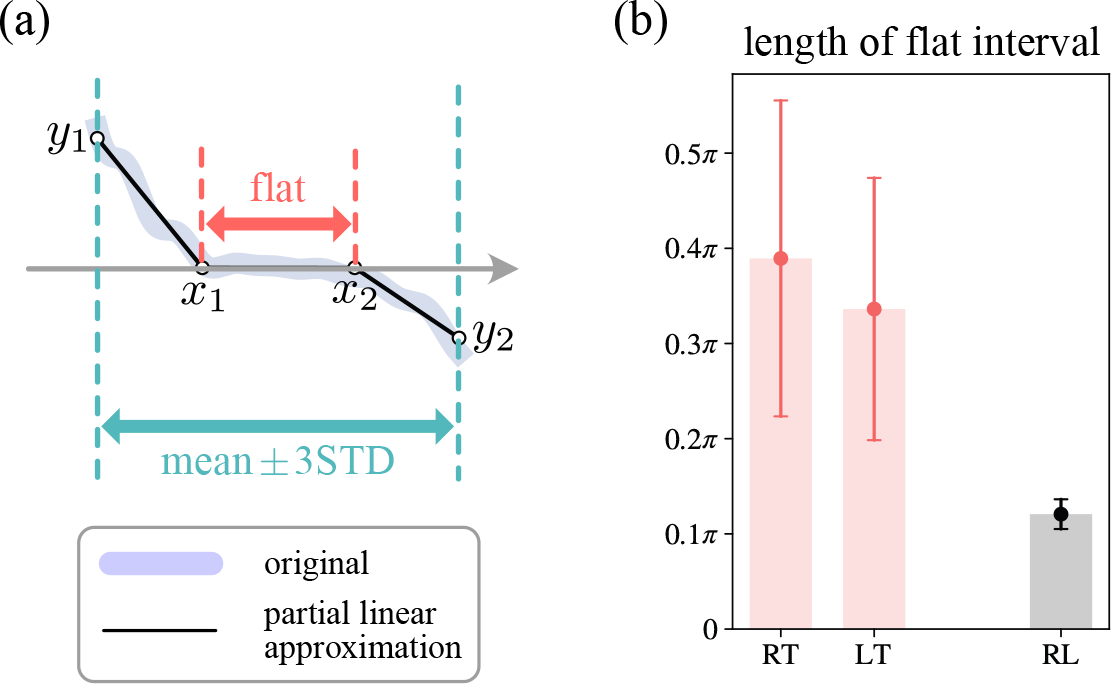
Approximation of piecewise linear function. (a) Four parameters used for the approximation. (b) The mean and standard deviation of the length of the flat interval. The label “RT” and “LT” mean the result of the *f*_RT_ and the *f*_LT_, respectively. The label “RL” means the result of the previous study (Arai et al., 2023).

Figure 5(b) presents the mean and standard deviation of the length of the flat intervals. In both the RT-model and LT-model, the flat intervals are longer than that of the RL-model and exhibit considerable individual differences.

## 5 Discussion

### 5.1 biological significance

In this study, we investigated the rhythm control of legs and the trunk during human gait using the phase reduction theory and Bayesian estimation methods. The motion of legs and trunk was modeled as coupled phase oscillators, and the model identification was performed using the time-series data. We found that the legs and trunk exhibited 1:2 synchronization, and the control of the relative phase was not strict. Similar results were observed in the rhythm control of both legs during gait, as reported in the previous study (Arai et al., 2023), suggesting that it may represent a typical control mechanism in the sagittal plane during walking.

The biological advantage of not strictly controlling the relative phase lies in the ability to conserve energy. However, a significant deviation from synchronous posture lead to deteriorated motor performance, necessitating the control actively. The weak stability of the relative phase observed in this study may align with such biological imperatives.

Moreover, intermittent control has been proposed in stabilizing upright posture (Asai et al., 2009; Suzuki et al., 2020), showing similarities to the results of this study.

### 5.2 mlimitations

The greater variability of the relative phase between the legs and the trunk, compared to that between both legs, may stem from the higher degree of freedom in the trunk. The trunk is flexible, particularly bending significantly in the sagittal plane. Consequently, the line segment used to reconstruct trunk phase in this study may not appropriately represent trunk behavior, suggesting the need for phase reconstruction with more information. This is left for future work.

While we considered only the interaction between single leg and trunk, it should be acknowledged that both legs and the trunk interact. Because identifying models for multi-body systems requires more data, we would like to measure a greater volume of time-series data and clarify the coordination between both legs and the trunk. We would also like to clarify the coordination involving both arms.

Finally, it is important to note that the participants in this study were limited to healthy men. Results may not necessarily extend to women, elderly people, or those with walking impairments.

### 5.3 does big data change neuroscience?

In recent years, there has been an increase in research exploring trends through big data (Dipietro et al., 2023). However, merely possessing a large volume of data does not necessarily yield meaningful results.

To grasp overall trends of big data, its visualization is effective, which requires some dimensionality reduction. In this study, we focused on the phase, one dimensional variable representing the rhythm of the system, which we reconstruct from the high dimensional time-series data. As a result, we were able to directly observe the synchronization between the legs and the trunk (Figure 3). Thus, variable selection tailored to the phenomenon of interest is crucial.

Furthermore, some mathematical model is indispensable for interpreting big data. In this study, we assumed a simple model of coupled phase oscillators and successfully extracted rhythm characteristics.

Thus, mapping big data to lower dimensions, it becomes possible to capture its macroscopic features, and interpreting it becomes feasible through estimating mathematical models from big data. We believe that big data will change neuroscience.

## Acknowledgement

This work was supported in part by MEXT KAKENHI Grant Number JP23H04467; JSPS KAKENHI Grant Number JP20K20520; and JST FOREST Program Grant Number JPMJFR2021.

## Notes

### Competing Interest Statement

The authors have declared no competing interest.

